# A generic systems-theoretic approach to identify biological networks capable of adaptation

**DOI:** 10.1101/2021.05.27.445914

**Authors:** Priyan Bhattacharya, Karthik Raman, Arun K. Tangirala

## Abstract

Constructing biological networks capable of performing specific biological functionalities has been of sustained interest in synthetic biology. Adaptation is one such ubiquitous functional property, which enables every living organism to sense a change in its surroundings and return to its operating condition prior to the disturbance. In this paper, we present a generic systems theory-driven method for designing adaptive protein networks. First, we translate the necessary qualitative conditions for adaptation to mathematical constraints using the language of systems theory, which we then map back as ‘design requirements’ for the underlying networks. We go on to prove that a protein network with different input–output nodes (proteins) needs to be at least of third-order in order to provide adaptation. Next, we show that the necessary design principles obtained for a three-node network in adaptation consist of negative feedback or a feed-forward realization. Interestingly, the design principles obtained by the proposed method remain the same for a network of arbitrary size and connectivity. Finally, we prove that the motifs discovered for adaptation are non-retroactive for a canonical downstream connection. This result explains how complex biological networks achieve robustness while keeping the core motifs unchanged in the context of a particular functionality. We corroborate our theoretical results with detailed and thorough numerical simulations. Overall, our results present a generic, systematic and robust framework for designing various kinds of biological networks.

## 1 Introduction

All living cells display a remarkable array of functions, which can be perceived as the response of a complex, multi-level biological network at a systems level. These complex networks are comprised of a variety of components— biological macro molecules—wired together in exquisite fashion. How the wiring of these components affects system function has been a classic subject of research over the last two decades. A variety of mathematical modeling techniques have been employed to model and predict the function of various biological networks [1–3]. Beyond mathematical modeling, systems theory has been particularly useful to understand and characterize various biological systems [4]. Graph-theoretic tools have also found applications in analyzing and understanding biological networks as functional modules [5–9]. Notably, it has been seen that the design principles, for any given biological response, are relatively conserved across organisms [10]. For instance, it is well-known that the adaptation (definition to be reviewed shortly) involved in performing bacterial chemotaxis in *E. coli* employs negative feedback. Similarly, an adaptive homeostasis network in higher organisms [11] also uses a negative feedback control strategy [12], suggesting relative independence of design principle from the particularities of the rate dynamics for different biological networks. This observation serves as an essential motivating factor behind the search for minimum networks capable of achieving a given biological functionality.

Besides adaptation, several studies have focused on understanding the emergence of functionalities such as oscillation, toggle switches, and determining the underlying circuitry [13–16], employing methods ranging from brute force searches [16] and rule-based modelling [13] to control-theoretic approaches [15]. Tyson *et al* (1974) conceived a two-protein negative feedback model with specific rate kinetics to prove the existence of an invariant Poincaré–Bendixson annulus which can lead to oscillation [13]. Li *et al* (2017) employed a brute force search across the topology–parameter space and concluded that incoherent self-loops and negative feedback provide robust oscillation in protein systems [14]. Sontag *et al* (2004) showed the necessity of positive feedback to attain a switch-like behavior which plays a crucial role in cell-fate decision making and quorum switching [17].

Adaptation is defined as the ability of the system output (O) to sense a change in the input (I) from the surrounding environment and revert to its pre-stimulus operating state. From the widely discussed bacterial chemotaxis [12], to the regulation of temperature in a volatile environment, or homeostasis, adaptation is believed to have played a pivotal role in evolution [18]. Typically, adaptation is characterized by two key quantities [10], precision and sensitivity. Precision is the ratio of relative changes of input and output and is quantified as

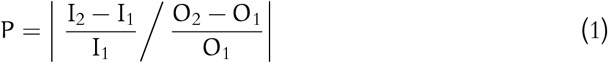

where, I_2_ is the new input, I_1_ is the initial input, O_2_ is the new output steady-state level, and O_1_ is the pre-stimulus output level. If O_2_ = O_1_, *i. e.* the system’s response returns to *exactly* the pre-stimulus level, the adaptation is known as *perfect adaptation*. On the other hand, sensitivity refers to the ratio to the relative difference between the peak value of the output (O_peak_) and the initial steady-state to that of the input:

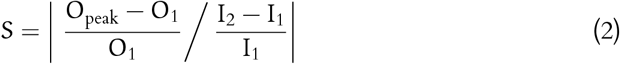

Previously, Tang and co-workers [10] investigated three-protein systems that were capable of perfect adaptation. A three-protein system, including self-loops, involves nine possible interactions, each of which can be positive (activating), or negative (repressing), or absent, resulting in a large number (3^9^ = 19, 683) of possible network structures or topologies. A brute force study of all the possible structures was carried out assuming Michaelis–Menten kinetics for the protein interactions. Each topology was examined for 10, 000 different sets of parameters leading to over 1.6 × 10^8^ simulations. The topology– parameter combinations that provided precision and sensitivity more than 10 and 1 respectively were considered capable of adaptation. Their study showed that only 395 topologies could perform robust adaptation. Surprisingly, all of the admissible structures had either negative feedback associated with a buffer species or incoherency in the input node’s effects on the output via two different paths. Later, other systems such as voltagegated sodium channels and gene regulatory networks were observed to exhibit adaptation as well. Notably, all the deduced structures employed negative feedbacks [19–21]. Sontag *et al* (2003) argued from an internal model principle perspective that attainment of adaptation with respect to a step-type disturbance requires an integrator within the system [22]. This, if used for a three-protein network, produces topologies similar to the 395 topologies discussed above.

Further, others have suggested specific control strategies like integral feedback to be capable of producing adaptation for a small network (containing three nodes) from an internal model principle and transfer function point of view [23–28]. Sontag *et al* [29] suggested the supremacy of negative feedback loops over the incoherent feed-forward structures in the context of providing adaptation to periodic responses with varying duration for small scale network structures. We have previously employed a transfer function approach to deduce the design principles for adaptation in a three-node network [26]. The main arguments were that the condition for perfect adaptation requires the transfer function of the system to be stable and contain a zero on the origin. Recently, Araujo and Liotta [30] developed a graph-theoretic method arguing that the feedback and feed-forward strategies are the only two ways of providing adaptation for networks with an arbitrary number of nodes and edges.

The present work provides a generic control-theoretic method using a state-space framework and shows that either negative feedback and incoherent feed-forward loop are necessary conditions for adaptation. In this sense, the necessary requirements on the network structure obtained through this work are more accurate and stronger than the previous studies. Our entire algorithm is independent of the kinetics, barring some minimal assumptions. This approach is in agreement with, and a generalization of the findings from previous studies [5, 10], which have argued that the structure of the network plays a determining role for the governing functionality.

The proposed approach enables us to identify all possible control strategies without resorting to a computationally demanding brute-force approach that can achieve perfect adaptation. We argue that the presence of either *negative* feedback or incoherent feed-forward loop are the only two ways to achieve adaptation. Besides, the proposed work also discusses the cases for adaptation for a staircase-type disturbance. We argue that a system that meets perfect adaptation is also capable of producing peak response in the minimum time. Further, we propose that the adaptive behaviour is invariant to a canonical downstream connection, which in turn shows the context-independence property of the adaptive networks, as opposed to oscillatory networks [31].

The rest of this article is organized as follows. The Methodology Section presents key concepts leading to the proposed algorithm, where the conditions for perfect adaptation are translated into certain equality constraints on the parameters of systems theory. The question of minimum peak response time is also addressed in this section. In the Application Section, the postulated mathematical conditions are used to identify the potential network structures of any size for adaptation The particular case of retroactivity in adaptation is also explained in the proposed mathematical framework of control theory. The final Discussion Section places the results along with the simulation studies in perspective.

## 2 Methodology

In this section, we outline a generic framework to deduce network structures capable of adaptation. First, we derive the mathematical requirements for the condition of adaptation using linear systems theory. Using these conditions, we first discover the motifs for adaptation by networks with a minimum number of nodes and edges. These conditions are further scaled-up to determine the necessary conditions for adaptation in networks of larger sizes, with arbitrary numbers of nodes and edges.

### 2.1 Linearisation of the rate reactions

Working in the linear domain allows us to utilize the wealth of linear systems theory. Given an enzymatic reaction network, the rate equations for the nodes, *i. e.* enzyme concentrations (**x**) can be written as

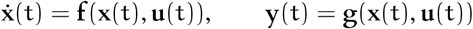

where **x**(t), **u**(t) and **y**(t) are the states, inputs or known disturbances and output, respectively. For this set-up, the linearized state-space model is

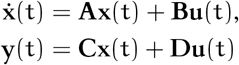

where **A**, **B**, **C** and **D** are obtained as the Jacobians of **f**(**x**, **u**) and **g**(**x**, **u**) with respect to the **x** and **u**, respectively. The corresponding transfer function can be written as

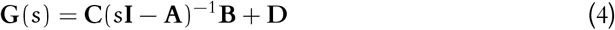

For the problem under consideration, the output and input are scalar variables. However, the obtained results apply to multiple-input, multiple-output (MIMO) systems. Indeed, a linearized model around a steady state does not always capture the non-linear dynamics accurately. However, since adaptation is a stable (convergent) response, according to the Hartman–Grobman theorem [32], the conditions obtained for adaptation using linear time-invariant (LTI) systems theory serve as sufficient conditions for the same even in non-linear systems.

### 2.2 Conditions for perfect adaptation

Perfect adaptation, as defined above, refers to a system that should be sensitive to changes in the input in its transient phase and be able to drive the response to its previous steady-state value. These conditions can be translated to restrictions on the state space matrices using LTI systems theory as (i) a non-zero peak value and (ii) a zero final value of the output.

The condition of non-zero peak value translates to a non-zero value of the sensitivity. This condition can be attained by making the output mode of the system *controllable* by the environmental disturbance. This can in turn be guaranteed, if the Kalman controllability matrix, Γ_c_, is full row rank, *i. e.*, for an N-dimensional state space with a single input,

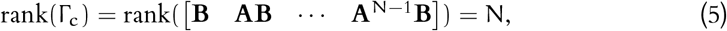

Since the system of rate equations are linearized around a stable fixed point, the initial value of the deviated output (deviation from the stable point) of the linearized system should be zero. These conditions, along with the assumption of linear, exponential stability (matrix **A** is Hurwitz), can be mapped onto the parameters of an LTI system for a step-change in the external environment, u(t), as

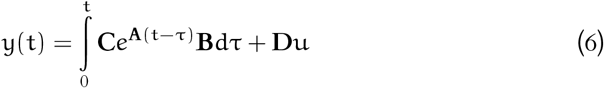

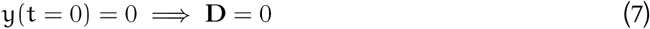

Using (6), the condition for zero final value can be obtained as

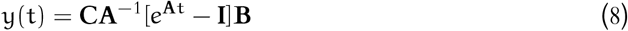

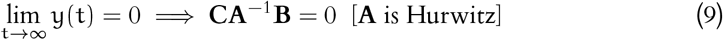

It is to be noted that the zero final value condition may not be achieved in several practical scenarios, leading to *imperfect* adaptation [33]. However, we shall limit this discussion to perfect adaptation. In this sense, adaptation and perfect adaptation shall be used interchangeably from here on.

Although (9) and (5) constitute the main checkpoints for adaptation, several other additional constraints, such as minimum peak time and minimum settling time can play a crucial role in sensing the change in the external disturbance and promptly acting to reject it. We argue below in Theorem 1 that the peak time for a system is minimum if the condition of zero final value is satisfied:

#### 1. Theorem

*For a set* 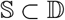 *(where* 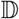 *is the ring of all causal transfer functions with real poles) consisting of stable, minimum phase transfer functions with the same set of poles and differing by a single zero position with each other, the transfer function with zero final value has the minimum peak time.*

*Proof.* To establish this fact, let us assume a proper LTI system G(s) and another system H(s) with same singularities (all real), except a zero at the origin. Assume y_1_(t) (Y_1_(s)), y_2_(t) (Y_2_(s)) and 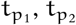 to be the step responses and the peak times for G(s) and H(s), respectively.

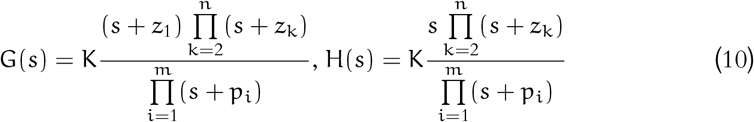

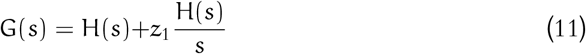

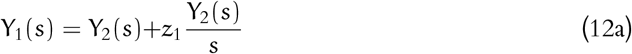

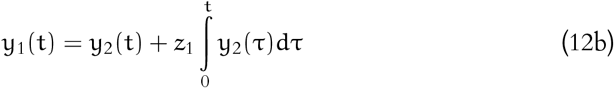

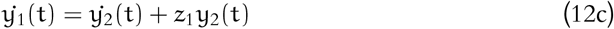

Setting 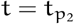,

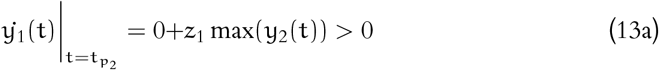

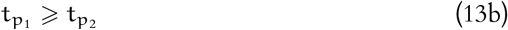

The equality in (13b) holds only when G(s) = H(s), *i. e.* g(t) shows perfect adaptation.

The above result can be extended in the case of damped oscillatory systems as well. From (12a), it can be seen that

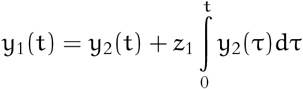

The peak time for y_2_(t) is always less than or equal to that of its integral 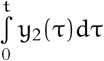 therefore their combination y_1_(t) has a peak time always greater than or equal to that of y_2_(t). Therefore, Theorem 1 implies that perfect (theoretically infinite) precision also ensures minimum peak time if the positions of the poles and the rest of the zeros are unchanged. The minimum settling time requirement involves calculating time constants, which for a large network can be obtained through a simulation study across different sets of time constants while retaining the property of zero final gain ((5)) to ensure perfect adaptation. To summarize, the conditions for adaptation derived above can be broadly divided into two sets. The first set of conditions ((9)) take care of the criteria for infinite precision, which includes the stability of the system matrix **A** and zero final gain of the step input. The second set ((5)) ensures non-zero sensitivity. This includes the controllability condition. Moreover, for a given network with a specific input–output configuration (*i. e.* with given **B** and **C** matrix), if the attainment of one set of conditions *ipso facto* violates other, then the network with the given input–output node cannot provide adaptation (see Fig. 1). In that case, a modification of the output node (since the input node is fixed for most of the practical cases) may resolve the problem.

**Fig 1.**
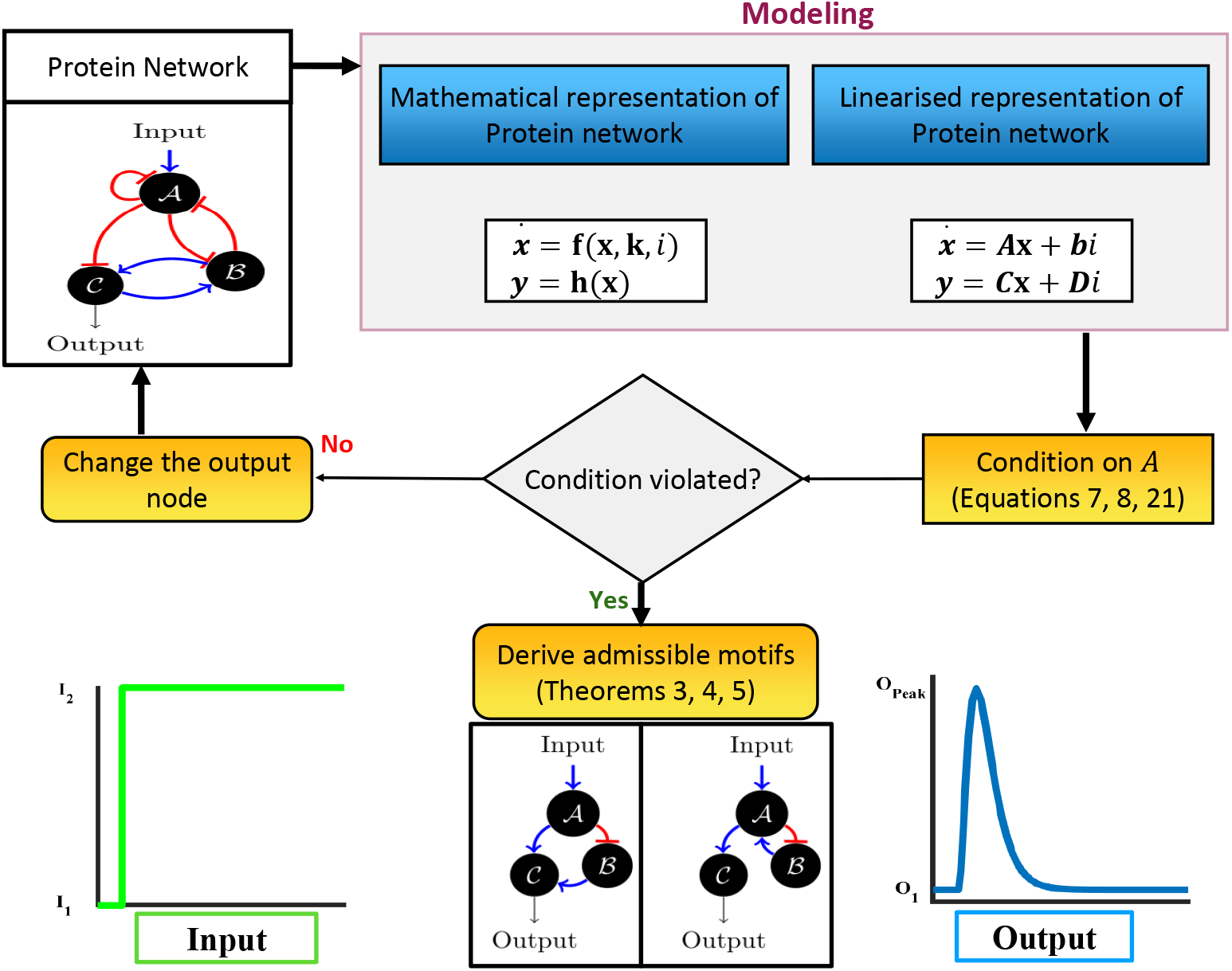
Workflow of the proposed methodology. Any given protein network is first linearized, and the conditions on the **A** matrix are investigated, to ultimately derive admissible motifs for the desired functionality.

## 3 Results

We demonstrate the capability of the methodology we developed above by applying it to protein enzymatic networks, where each node is a protein, and an edge represents either of the following:

1. *Activation*: a protein 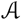 is said to activate 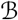 when 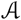 acts as a transcription factor that binds the active site of the promoter of 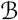 to aggravate the transcription process for the synthesis of 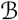.
2. *Repression*: similarly, if 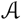 acts as a transcription factor to reduce the transcription rate of mRNA, which translates to 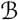.

For a network containing N nodes, there are 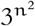 numbers of possible network structures. The generalized state equations for an N-node network can be written by considering the normalized concentration of each protein as states:

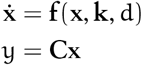

where, **x** ∈ ℝ^N^ and **k** ∈ ℝ^p^ are the states and the parameters associated with the rate equations. In passing, it may be noted that in the presence of any algebraic constraints on the states (e.g. due to conservation laws), an N-node network corresponds to a reduced-order dynamical system. In the single disturbance case, d is referred to as the disturbance variable. The protein that receives the external disturbance directly is considered as the input node. The concentration of the N^th^ node is taken as the output.

### 3.1 Two node networks– are they capable of adaptation?

From a systems theoretic viewpoint, the step response of a first-order system is always a monotone which is not the case with adaptation. Therefore, the possibility of providing adaptation for any single protein can be safely ruled out. The immediate next case of N = 2 can be investigated. Implementing the aforementioned approach (Fig. 1) reveals that two protein networks with different input and output nodes are unable to provide adaptation. However, two-node networks with the same input and output nodes can perform adaptation (see *SI Methods*) as shown in 2.

It is important to note that the system matrix **A** for an N-node system linearized around a stable operating point carries not only the necessary information about the structure of the network but also the type of each edge, *i. e.* activation or repression. For instance, if 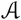 represses 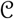, the element in the associated **A** matrix that corresponds to this edge turns out to be negative. This implies that f_ij_(**x**) (for activation) or −f_ij_(**x**) (for repression) is a class 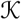 (*i. e.* monotone within a finite open interval in the domain, and passes through the origin) function with respect to x_j_, ∀j ≠ i. Intuitively, **A** matrix acts as a variant of the incidence matrix for the graphical network, with the diagonals being the exceptions. It is possible to have a negative or non-positive value of the diagonal element, albeit in the presence of a self-activation loop (refer to *SI Methods*). These inherent properties of the biological systems’ rate dynamics perform an instrumental role in maintaining the structural determinism property of adaptation.

Interestingly, there exists a class of biological networks that provide adaptation for a single step input but do not respond to subsequent perturbations [21]. This is defined as the *toilet flush phenomenon* (Fig. 2). Friedlander *et al* (2011) and Goh *et al* (2013) showed that this phenomenon occurs in a three node network with an equality constraint stemming out from a conservation law thereby reducing the effective number of state variables to two [19, 34]. In this regard, the aforementioned algorithm provides a great systems-theoretic perspective to explain and design such networks. If the time difference between two successive step perturbations is large enough (compared to the system’s settling time), then the condition for adaptation in this case is the same as that for a single step. Along with this, it is to be observed that with each step perturbation, the steady-state values of the system changes (note that the adaptation property guarantees the invariance of the steady-state of the output state only), which leads to a different linearized model. If the modified linearized model remains controllable and the general condition of adaptation is satisfied, the system provides adaptation for staircase input (Refer to SI Methods for a detailed discussion).

**Fig 2.**
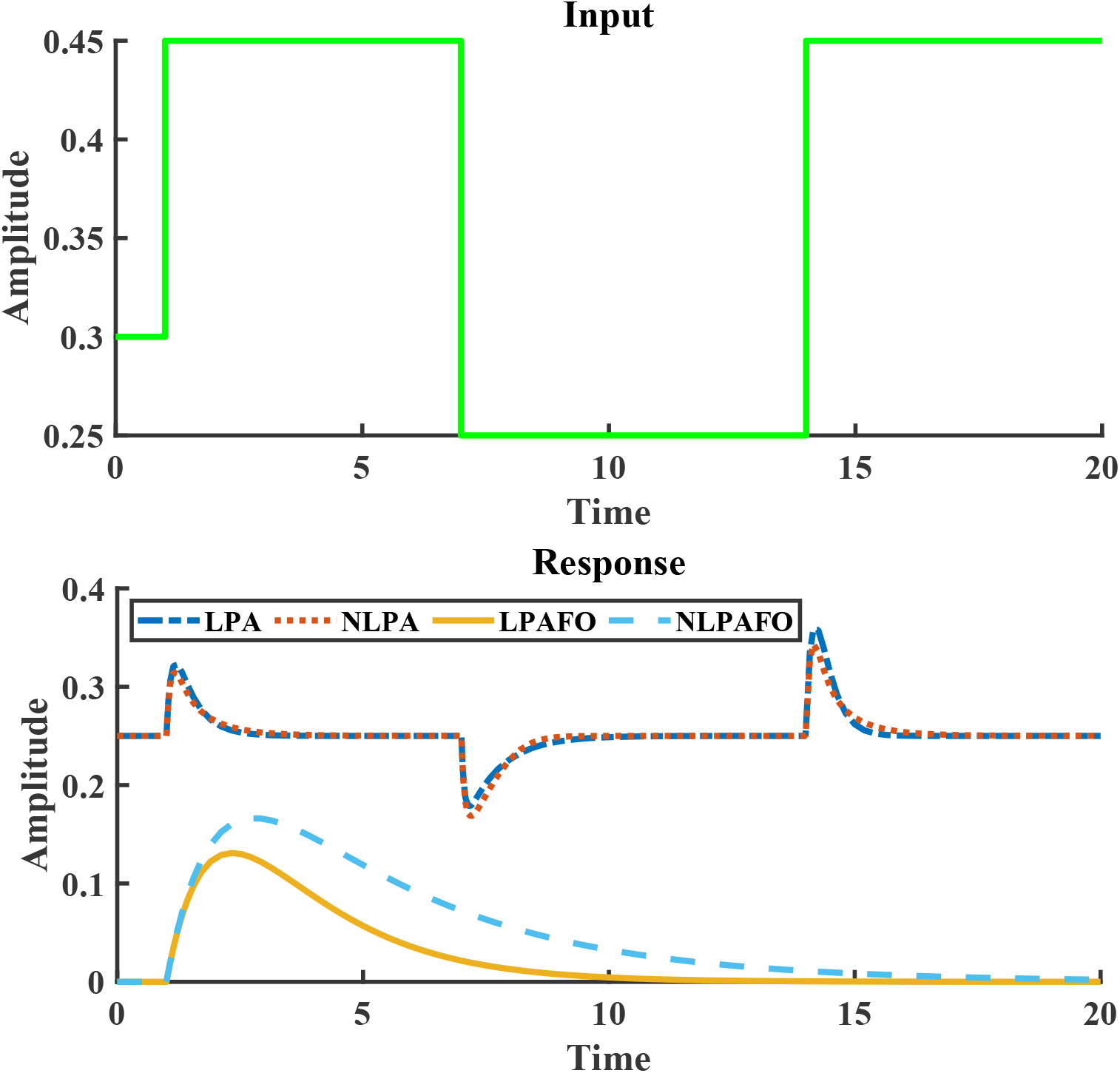
Two node networks capable of adaptation subject to staircase disturbance. The abbreviations LPA, NLPA, LPAFO, NLPAFO stand for linear perfect adaptation, nonlinear perfect adaptation, linear perfect adaptation for once and nonlinear perfect adaptation for once respectively.

### 3.2 Three-node networks with a maximum of three edges

The admissible network structures obtained from the analysis of the two-node enzymatic networks exclude the possibility of network structures that can provide adaptation with different input-output nodes. Therefore, it is important to identify a control strategy– perhaps the inclusion of an additional controlling node–that can bring adaptation to the two-node protein system with different input–output nodes.

From the perspective of a control-theoretic framework, the functionality of adaptation can be thought of as a regulation problem. Considering the biological feasibility and the network with only one external disturbance input 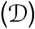, we propose a feedback control scheme where another protein 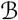 can act as a controller node. If the concentration of 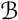 is u, the controller dynamics can be written as

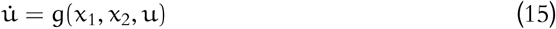

We adopted g : ℝ^3^ → ℝ as a linear function of the states and the control input.

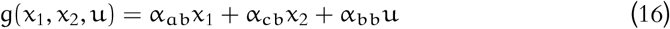

The parameters such as α_ab_ and α_cb_ govern the strength and type (repression or activation) of the edges. From feedback control theory [32] if the open-loop system is fully controllable by u then consideration of u as a variant of dynamic state feedback control strategy does not alter the controllability of the system.

#### 3.2.1 Finding the admissible topologies

The closed system can be written as

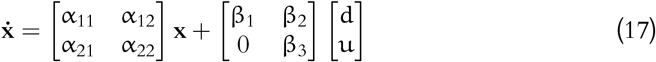

For the system to provide adaptation, x_2_ has to be controllable by the control input u. For the closed-loop system, the infinite precision condition for adaptation can be written as 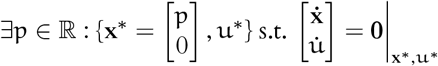

For the system with controller,

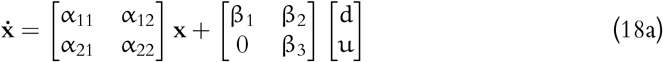

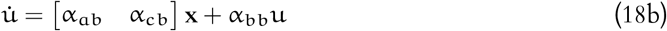

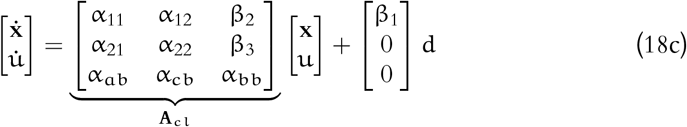

Using the condition for adaptation,

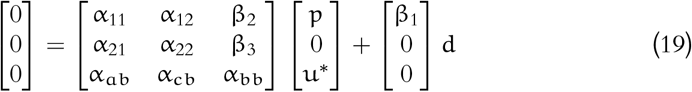

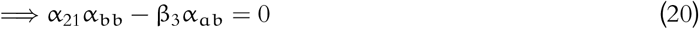

The condition α_21_α_bb_ − β_3_α_ab_ = 0 can be achieved in three scenarios:

1. All the terms are zero: this leads to singularity of **A**_cl_, and is hence not acceptable.
2. α_21_α_bb_ = β_3_α_ab_ = 0: this is feasible. Interestingly, if α_21_ = 0, the state x_2_ becomes unobservable. Also, in order to attain the condition for adaptation, making α_21_ = 0 requires either (i) β_3_ to be zero, which in turn, results making x_2_ an uncontrollable mode with respect to u or alpha_ab_ = 0 leading to uncontrollability with respect to i.
3. α_21_α_bb_ = β_3_α_ab_ ≠ 0: this is acceptable as long as **A**_cl_ is Hurwitz.

Combining each of the feasible possibilities with the infinite precision condition for adaptation, we arrive at a superset of admissible motifs from the above possibilities. As it can be seen from Table 1, the first three network motifs involve negative feedback engaging node 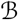. This type of network can be termed as *negative feedback loop with a buffer node (NFBLB)*. Since NFBLB involves negative feedback, the corresponding response becomes damped oscillatory for most of the cases. However, as long as the adaptation criterion is satisfied, the output after a damped oscillatory transient response goes back to its initial steady state.

**Table 1.**
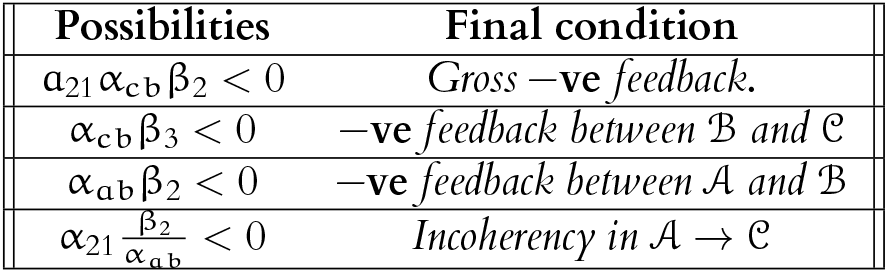
Possible motifs for adaptation

| Possibilities | Final condition |
| --- | --- |
| $\alpha_{21}\alpha_{cb}\beta_2 < 0$ | Gross $-\mathbf{ve}$ feedback. |
| $\alpha_{cb}\beta_3 < 0$ | $-\mathbf{ve}$ feedback between $\mathcal{B}$ and $\mathcal{C}$ |
| $\alpha_{ab}\beta_2 < 0$ | $-\mathbf{ve}$ feedback between $\mathcal{A}$ and $\mathcal{B}$ |
| $\alpha_{21}\frac{\beta_2}{\alpha_{ab}} < 0$ | Incoherency in $\mathcal{A} \rightarrow \mathcal{C}$ |

The remaining motif carries an incoherency between the two forward paths (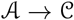 and 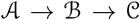) from 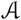 to 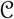. This is precisely the reason it is called *incoherent feed-forward loop with proportioner node (IFFLP).* Owing to the structure of IFFLP, the underlying system matrix **A** for IFFLP will always have real eigenvalues, thereby eliminating the possibility of oscillatory transients (Fig. 3).

**Fig 3.**
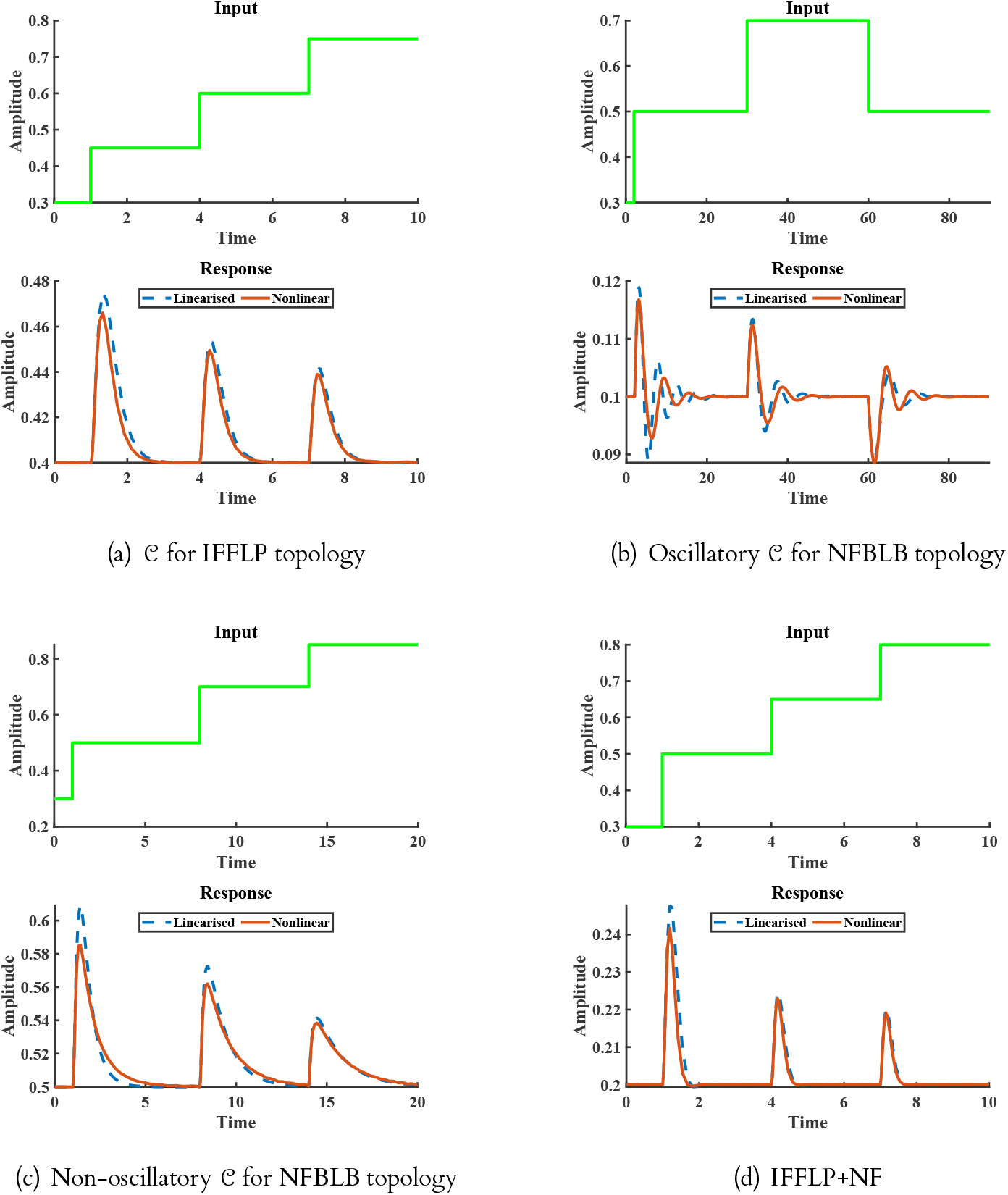
(a) shows the response of the output node for a three-node IFFLP topology. (b) shows the same for a three-node NFBLB. The oscillatory behavior can be attributed to the complex eigenvalues of the **A**. Similarly, (c) shows a non-oscillatory response of an NFBLB motif. (d) is the response of the output node of a network containing both the admissible network structure *i. e.* incoherent feedforward path and negative feedback.

### 3.3 Finding all possible three-node motifs capable of adaptation

After finding the minimal network structures—minimal in terms of edges and number of nodes—we extend the above method to find the necessary topological properties, *i. e.* the existence of feedback or feed-forward configurations without any restriction on the number of edges, for the three-node network.

*Remark* 1: For any three-node network, the corresponding system matrix can be written as

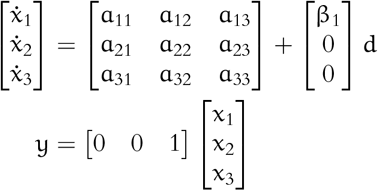

For adaptation, 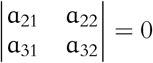 and **A** has to be Hurwitz.

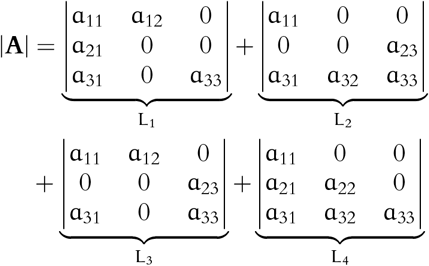

As it can be seen, the determinant of **A** can always be written as a combination of determinants of elementary topologies containing exactly 3 edges. For **A** to be Hurwitz, |**A**| has to be negative, *i. e.* at least the determinant of any one of these four matrices has to be negative. If any of the first three terms (L_1_, L_2_, L_3_) is negative, it indicates negative feedback. Note the condition 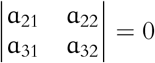 is ‘structurally’ satisfied for L_1_, L_2_, L_3_ but in the case of L_4_ it has to be satisfied by the parameters. If L_4_ is the only negative term, then there exists an incoherent feed-forward loop in the network. Similarly, multiple negative terms represent the presence of both types of motifs. This implies that for any three-node network capable of adaptation with arbitrary edges, the presence of negative feedback or incoherent feed-forward loop is a *necessary condition*.

Since the negative determinant for **A** ∈ ℝ^3×3^ is a weaker condition for stability compared to that of **A** being Hurwitz, the presence of either or both incoherent feed-forward and negative feedback loops is only a necessary but not sufficient condition for adaptation.

Figure 3 depicts the response of different admissible three-node networks to identical disturbance input. Similar to 2 the signals expressed in lines and dots refer to the responses of the non-linear rate dynamics and corresponding linearised counterparts for the corresponding network structure, respectively. In both the cases a variant of Michaelis Menten kinetics is considered for simulation. It can be inferred from figure 3(a) that IFFLP always produces hyperbolic responses. The reason behind this can be traced to the spectrum of the underlying system matrix **A** ∈ ℝ^3×3^ in the linearised dynamics. Due to the absence of any loop in the network, the associated **A** matrix for a feedforward network is lower triangular, with the diagonals being the eigenvalues, thereby resulting in hyperbolic responses. Unlike IFFLP, NFBLB can potentially give rise to oscillatory responses along with perfect adaptation as shown in figure 3(b).

The above framework, developed for three-node networks, can be extended to larger networks with N–nodes and P–edges. As shown in the previous section, a three-node network comprising an input, output, and controller can provide adaptation. In this sense, an N (N ⩾ 3) node network can be thought of as the closed-loop system incorporating I/O nodes along with the controller network comprising of the remaining N − 2 nodes. At first, we derive the admissible elementary N–node network structures *i.e.* networks that contain at most N–edges and can provide perfect adaptation. We then use these results to establish the necessary structural conditions for perfect adaptation in case of any N–node network.

### 3.4 Condition on minimum number of edges in an N—node network for adaptation

In the following theorem, we first derive the lower limit on the number of edges required for an N−node network to provide perfect adaptation.

#### 2. Theorem

*For a network with* N ⩾ 3 *nodes, at least* N *edges are required to provide perfect adaptation.*

*Proof.* It has already been established that in the case of biochemical networks, the system matrix **A** for the *linearized* dynamics serves as the digraph generating matrix. Let us assume that the above statement in the theorem is wrong *i.e.* there exists an N− node, N − 1 edge network that can achieve adaptation. For an N-node, N−1-edge network to show adaptation, it has to satisfy (i) the controllability condition and (ii) infinite precision condition. The mathematical expression for the second has already been derived in (9). However, here we modify the equation for convenience.

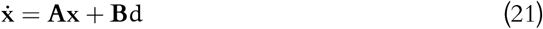

where, **x** = [x_1_ x_2_ ⋯ x_N_]^τ^ ∈ ℝ^N^ is the state vector with each element (x_i_) representing the concentration of each node (i^th^ node) and **B** = [β 0 ⋯]^τ^ ∈ ℝ^n×1^. Let the output be concentration of the K^th^ node and input be applied on the first node. This implies that the steady state concentration 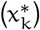 of the output node is zero in linearised representation. At the steady state,

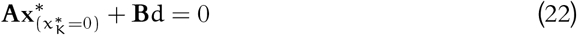

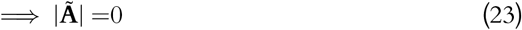

where, 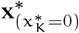 is the steady state solution to (21) with the K^th^ component being zero and 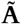 is the minor of the component representing the edge from the output to the input node.

From elementary network theory, it can be said that it is always possible to design a controllable network with N nodes with N − 1 edges if and only if there is no feedback loop. Since the possibility of an isolated node is eliminated, the only feasible structure for a N-node, N − 1 edge is a feed-forward network (N-node networks with a lower number of edges are eliminated for the same reason). Further, since the number of edges is N − 1, no node can have more than one incoming or outgoing edge. In order to satisfy the controllability condition, it requires at least one forward path from input node to the output. In the case of an N-node network with N − 1 edges and no isolated nodes, there can exist one forward path from the input to output node maximum.

The second condition, *i.e.*, the infinite precision condition requires the minor of the component of **A** matrix that represents a direct edge from the input to output node be zero. For any digraph matrix in ℝ^N×N^, every term in the determinant expression contains N! terms, each a product of N−tuples chosen from the matrix. Further, from combinatorial matrix theory, each of these N−tuples can be expressed as a multiplicative combination of the matrix elements that map to existing loops of the network and the diagonal elements. According to this result, each term in the minor of **A**_1K_ has to contain at least one forward path from the first to the K^th^ node. Since in the case of N − 1 edge networks, there can only be one forward path possible, the minor of **A**_1K_ is a singleton set. Thus fulfilling the infinite precision condition in this scenario amounts to deleting the only forward path from the input to the output node rendering the system uncontrollable (See *SI Appendix*). On the other hand, it has been observed that when N = 3, the number of edges required to produce adaptation is also three (more generic demonstration of constructing N-edge N-node motifs that can achieve adaptation is provided in the supplementary information). By virtue of the foregoing discussion, we conclude that the minimum number of edges required for adaptation is N.

### 3.5 Feedforward networks are adaptive only when incoherent

We are now ready to present below the most essential and generic results emanating from this work. According to Theorem 2, it requires at least N edges for any N−node network to provide adaptation. It can also be shown that there exist only two principal means to satisfy (23) for any elementary N−node network(refer to SI Methods). The admissible elementary network structures can be divided into two further categories i) network without and ii) with loops. In the first scenario, we argue in the following theorem that the existence of at least two opposing feed-forward paths is a necessary condition for adaptation.

#### 3. Theorem

*For an* N−*node network without any loop, the only way to provide perfect adaptation is to have at least a pair of feed-forward paths from the input to output loop with opposing effects.*

*Proof.* Let us consider the concentration of the k^th^ node as the output variable. It can be shown that for the output variable to adapt to disturbances, k has to be greater than two (refer to *SI Methods*). Given an N−node, controllable network structure with no loops, it is always possible to order the nodes so that the resultant digraph matrix is lower diagonal. Since the system matrix **A** is equivalent to the digraph matrix, it shall also inherit the lower diagonal structure.

Assuming k > 2, for the output node of the network structure to provide adaptation, it has to satisfy the i) controllability ((5)) and ii)infinite precision (Eqs. 9, 23) conditions. It can be shown that a feed-forward network is always controllable (refer to *SI Methods*). Also, the lower diagonal property of **A** guarantees the stability of the system, given the diagonals are strictly negative.

The infinite precision condition in (23), requires the minor of the component **A**_1k_ (Denote it as 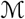) to be zero. From combinatorial matrix theory [35], it can be stated that except the product of all the diagonal elements, every other term in the determinant expression of a digraph matrix maps to the product of the diagonals and the loops. Using this result, it can be claimed that each of the (N − 1)! terms obtained through multiplying **A**_1k_ with its minor in the determinant expression of **A** is composed of products of the loops and diagonal elements. Also, each of these terms must contain exactly one loop that involves the edge from k^th^ to the first (input) node. Therefore, the corresponding terms in the minor of **A**_1k_ should contain exactly one possible forward path from the input node to the output node along with other possible loop or diagonal elements or both.

Since there are no loops in the feed-forward network structure, every term in the minor expression contains exactly one forward path and diagonal elements. Let us define the set 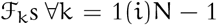 where each element in 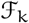 contains the product of the elements in the **A** matrix that represents a forward path with k edges and N − 1 − k diagonals with no common indices with the former. Consequently, the minor expression can be written as

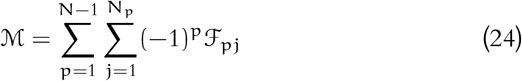

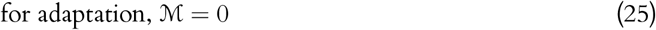

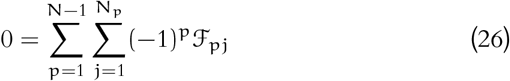

where, N_p_ is the cardinality of the set 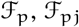 is the j^th^ element of 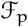. If 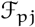 has a forward path f_pj_ and the product of the diagonals as D_pj_ the associated cumulative sign (S_p_) of 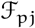 in the minor expression can be written as

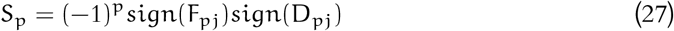

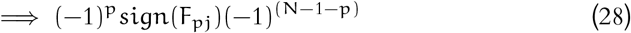

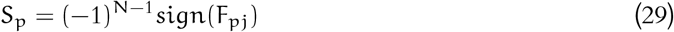

It is evident from (29), S_p_ is independent of p but a function of the effective sign of the forward path. For (26) to hold, there should be at least one pair with mutually opposed cumulative signs. This can only be possible if there exists at least one forward path with the effective sign being positive, and at least one of the remaining forward paths has to be of the effective sign negative.

### 3.6 Conditions on elementary networks with loops for adaptation

In the second case (N−node, N − edge networks with at least one loop), one of the possible network structures with N edges can be composed of two or multiple loops without any connecting edge and the common node. In the next theorem, we argue that this type of network cannot attain adaptation.

#### 4. Theorem

*An* N− *node network containing multiple loops with no common species and no edge connecting the loops cannot provide adaptation.*

*Proof.* As established in the methodology section, the underlying linearized dynamical system has to be controllable by the external disturbance input to perform adaptation. Now, for an N-edge network with L_m_ loops with no common nodes between them, the associated system matrix **A** can be written as

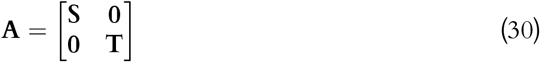

where, **s** consists of the loop element involving the input node x_1_ and **T** comprises elements representing all the remaining L_m_ − 1 loops. To avoid the trivial scenario, we consider that the output node is not involved in any loop with the input network. Let the loop involving the input node x_1_ involves n_1_ number of nodes then 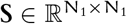 and 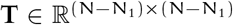. It is to be noted that if the output node is involved in a loop with the input node, then the effective network order reduces to N_1_. In order to avoid such trivial cases, we assume that the output is involved in any of the remaining L_m_ − 1 loops. Given 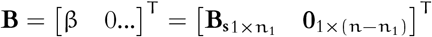 associated Kalman controllability matrix (**K**) for the pair (**A**, **B**) can be evaluated as

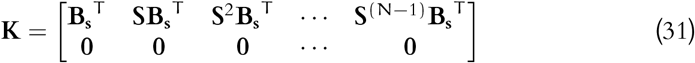

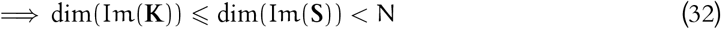

where, Im(·) denotes the column space of a matrix, and dim(·) calculates the dimension of a given vector space. From (32), it is clear that the Kalman rank condition is not satisfied in this case, leading to failure in achieving adaptation.

Therefore, the problem of uncontrollability discussed in Theorem 4 can be circumvented by placing at least one connecting edge between each loop with no common nodes. In that case, the question of stability has to be taken into consideration.

#### 1. Remark

*Along with the equality condition* (23), *the stability of the linearized system should also be guaranteed. Again, we impose a weaker condition of stability by invoking the sign of the determinant of* **A**. *If* **A** ∈ ℝ^N×N^ *is Hurwitz then*

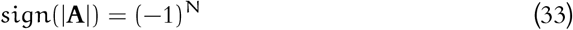

*It is to be noted that for any matrix* **A** ∈ ℝ^n×n^ *to be Hurwitz, it is required to satisfy exactly* N *number of conditions. The condition mentioned in* (33) *is one of them which is concerned with the product of the eigenvalues.*

With this stability criterion, it can be shown that specific network structures with positive feedback loops can not provide adaptation due to loss of stability.

#### 5. Theorem

*An* N-*controllable node network with multiple loops and no common nodes cannot provide adaptation if the effective signs of all the loops are positive.*

*Proof.* Let ℚ be the set containing all the controllable candidate motifs containing multiple loops with no common nodes but edges connecting each loop. Further, we assume that every node is involved in exactly one loop. Suppose, an element 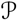 in ℚ consists of L_p_ number of loops. It is evident that for 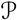 to be controllable, it has to contain N edges consumed by all the L_p_ loops along with minimum L_p_ − 1 edges connecting the loops summing up to N + L_p_ − 1 edges in total. In the present context, The minimal motifs can be thought of as the elements in ℚ which has N + L_p_ − 1 number of edges and L_p_ number of loops. Define the set Φ ⊂ ℚ consisting of all possible minimal motifs in ℚ.

In order for a minimal motif in Φ to provide adaptation, it must satisfy the adaptation condition (23). This can be achieved if and only if at least one of the diagonal elements of the **A** matrix is zero (refer to *SI*). Let us assume, that the 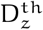 row of the **A** matrix contains the zero diagonal. Since none of the loops share any common node, the 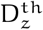 node must be associated with only one loop denoted by L_z_. Suppose L_z_ involves N_z_ number of nodes. The set 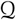 contains all the elements in **A** matrix that correspond the loop L_z_. Suppose 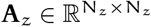 be the sub matrix of **A** that captures the connection patterns of all the N_z_ nodes involved in L_z_. Since each node is involved in only one loop the structure of the associated **A** matrix can be written as

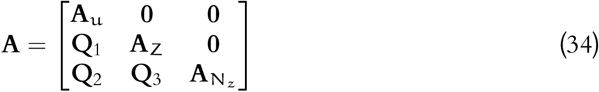

where, 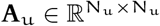 is the sub matrix that captures the upstream loops to L_z_ and 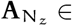 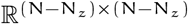 involves all the loops except the upstream loops and L_z_. The sub matrix **Q**_i_ captures the downward edges joining the loops. If the spectrum of **A**_z_, and 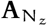 are 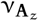 and 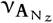 respectively then the spectrum of 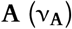 can be expressed as

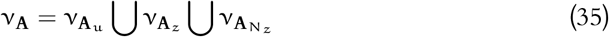

It is evident from equation (35) that for **A** to be Hurwitz, **A**_z_ has to be Hurwitz. Imposing the stability criterion as defined in the equation (33) on **A**_z_,

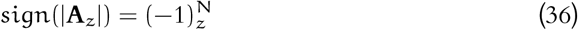

Since one of the diagonal components of **A**_z_ is zero, the determinant in this case is the product of all the elements mapping to all the edges involved in the L_z_. From combinatorial matrix theory [35], the sign assigned to a loop with N_z_ number of nodes in the determinant of a matrix can be written as 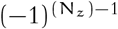.

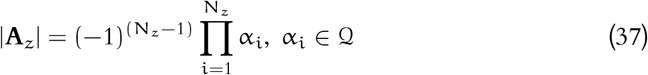

Therefore, using equation (36) we can say for **A** to be Hurwitz the following condition should hold

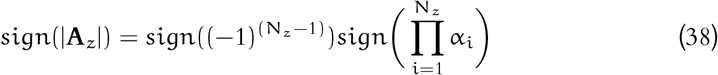

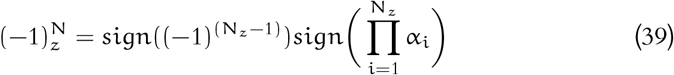

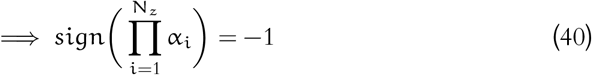

From (40), it is clear that, if the cumulative signs for all the loops of any candidate motif in Φ are positive then the resultant **A** becomes unstable, failing to provide adaptation.

### 3.7 Conditions for adaptation in larger networks

The above important results help us find out the necessary structural conditions for a network of N nodes and P edges, (∀N ⩾ 2, P ⩾ 2) to provide adaptation that are formalized below.

#### 6. Theorem

*Statement 1 serves as a necessary condition for Statement 2*

1. *There exists either negative feedback or incoherent feed-forward loop or both in the network*.
2. *The network attains perfect adaptation in the presence of a step type disturbance*.

*Proof.* From Theorem 2, consequently, if the rank of 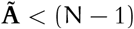 then it will lead **A** to be rank deficient, which in turn, violates the stability criterion.

The admissible N−node topologies that satisfy the adaptation condition (23) contain at most (N − 1)(N − 1)! number of elements in the determinant expression of the underlying linearised system matrix, **A**, Also, each of the (N − 1)(N − 1)! elements is the product of some N terms belonging to **A**. All the elements except the one that is the product of the diagonal contain at least two off-diagonals. Each term containing the product off-diagonal (and diagonals) terms of the system matrix refers to a loop when mapped back to the structure [35]. For instance, a term in the expression containing P off-diagonal elements (remaining N − p diagonals) can map to a loop engaging P nodes. This refers to a network containing a loop of P links and N − P forward paths (Theorem 2). So, using matrix theory, the prefix sign of each term in the determinant expression of any matrix can be determined by calculating the minimum number of exchanges needed to arrange them as products of diagonals.

For instance, an element with a loop with P-nodes, the remaining N − P are the diagonal elements. Now, for a P-node loop, the minimum number of exchanges necessary for arranging them as the product of diagonals can be easily obtained as P − 1. So, the stability condition for N-node network with a single loop with P nodes can be written as:

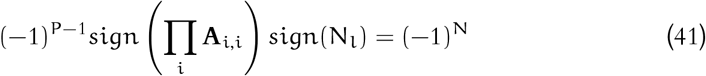

Using the assumption of all the diagonals to be default negative

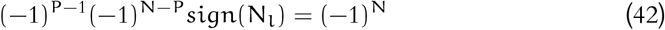

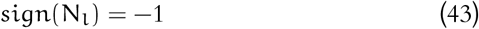

This concludes the presence of negative feedback loops as admissible elementary motifs for adaptation. As it can be seen from Theorem 3, the only term consisting of N diagonal elements can be written as the determinant of incoherent feed-forward loop motif as the associated **A** matrix for IFFLP is lower diagonal. Interestingly, it can also be shown that the elementary network structures *i.e.* the negative feedback loop and IFFLP satisfy the stronger Hurwiz stability criterion as well (Refer to SI).

So, for any network of arbitrary node N and arbitrary number of edges, if it attains adaptation, then its determinant can be written as the sum of the determinants of (N−1)(N−1)! numbers of elementary motifs. To satisfy the stability criterion, the determinant of at least one elementary motif should be of the sign (−1)^N^. This showcases the presence of either NFBLB or IFFLP as a universal, necessary condition for adaptation. The elements in the determinant expression mapping to multiple loops without common nodes can be concluded as incapable of adaptation using the second dependency. It can also be shown that these structures with a link between the nodes cannot provide adaptation if there exists no single negative feedback because it fails to satisfy the stability condition as at least one of the eigenvalues of the matrix becomes positive (refer to Theorem 5).

It is to be stressed that these structural conditions for adaptation only serve as necessary conditions for two reasons. Firstly, the sign of the determinant condition used here in (33) is only a weak (necessary) property of a stable system. Secondly, there are additional quantitative constraints that are to be satisfied by fine-tuning the parameters. For instance, in a three-protein system, the negative feedback requires α_bb_ = 0, which needs to be guaranteed by the parameters. Similarly, a three-protein network with incoherent feed-forward loop requires α_21_α_bb_ = β_3_α_ab_ ≠ 0 to be satisfied by the parameters.

Interestingly, it is found that adaptation is preserved against the connection with a downstream system (Fig. 4). The connection considered here is canonical, *i. e.* only the output node is connected with the downstream network.

**Fig 4.**
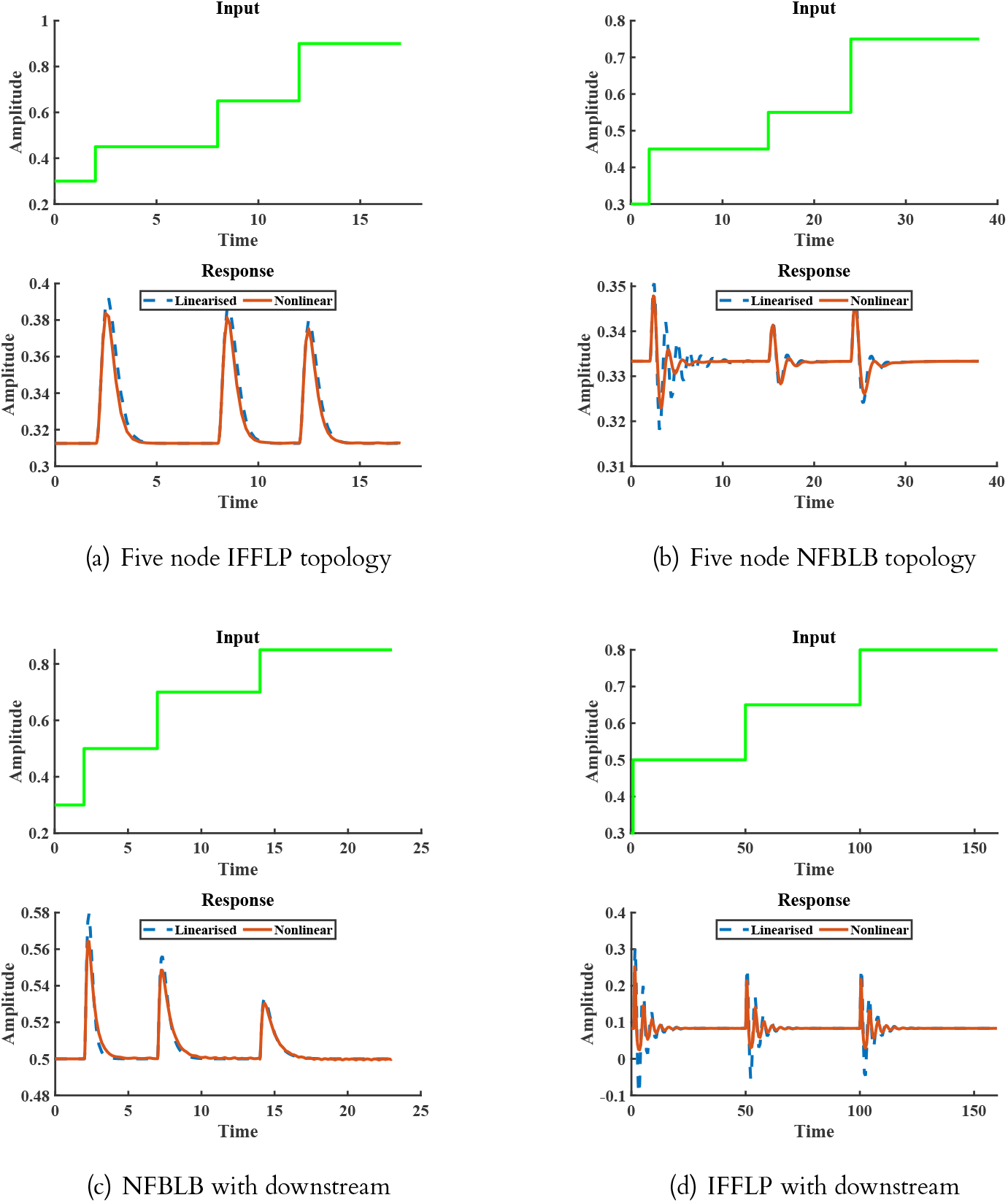
(a) shows the response of the output node for a five node IFFLP topology. (b) shows the same for a five node NFBLB with a hyperbolic response. The oscillatory behavior in (c) can be due to negative feedback, leading to complex eigenvalues of the underlying **A** matrix. (d) demonstrates the modular behavior of an NFBLB motif when connected to a downstream system. (e) is the response of the output node of an IFFLP network connected with a downstream system. Although the functionality of adaptation is not compromised, the oscillatory behavior is undoubtedly due to the negative feedback associated with the output of the IFFLP module and the downstream node.

#### 7. Lemma

*If the stability of the system is not altered, then the functionality of perfect adaptation for an upstream system does not get altered if the output node is connected with a downstream system.*

*Proof.* Given an upstream adaptive network containing N nodes and P edges, it is to be proved that the system preserves its functionality if it is connected with another arbitrarily connected network. Without any loss of generality, let us assume the 1st and the N^th^ nodes are the input and output nodes of the upstream network, respectively. The downstream system is connected in a feedback fashion with the output node.

Let the system matrices of the upstream and downstream networks be **A**_1_ ∈ ℝ^N×N^ and **A**_2_ ∈ ℝ^P×P^, respectively. As per the statement, the upstream system can provide adaptation, *i. e.* 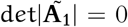, where 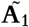 is the matrix associated with the minor of a_1N_. Due to the assumption of the structure, the modified system matrix **A**′ for the augmented system can be written as

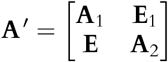

where, the elements of **E**_1_ ∈ ℝ^N×P^ are zero everywhere other than the N^th^ row. Similarly, the elements of **E** ∈ ℝ^P×N^ are zero everywhere other than the N^th^ column. For the combined system to produce adaptation, the minor of 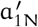 has to be zero. The matrix associated with the minor of 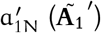 can be written as

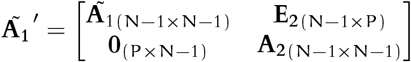

Since 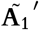 can be expressed as a block diagonal matrix with the lower non-square matrix being zero, the determinant is the product of the individual determinants of 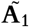 and **A**_2_. According to the assumption on the upstream system 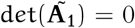, therefore the matrix 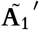 is singular. This, in turn, implies that the combined system can provide adaptation if the stability is not altered.

This is intuitively a well-expected result because, typically, adaptation networks are mounted on the big downstream network to provide robustness with respect to external disturbances, and the above lemma shows that the adaptation networks are not retroactive and context-dependent.

## 4 Discussion

Biological networks are complex yet well-coordinated and robust in nature. Although the form of the reaction dynamics underlying a network governs certain behaviors of the biological system, the major roles of controlling and coordinating different levels of hierarchy in the networks can be attributed to the very structure of the network. Previous research works have adopted one of a brute-force, graph-theoretic or a rule-based approach for identifying admissible structures for perfect adaptation. The nature of results obtained from these approaches are limited by the computational cost, inability to capture all necessary structures and/or the challenges in handling networks of arbitrary sizes. In this work, we appeal to the linear systems and control theory for obtaining formal and generalised results without being bounded by any of the aforementioned limitations.

Intuitively, it is apparent that for any (biological) system to exhibit adaptation, it should internally possess a feedback and / or feedforward configuration as mandated by control theory. However, deeper and concrete answers, especially on how such results scale-up with the size of network, inevitably call for a formal study. The primary questions that formed the basis of this work are (for perfect adaptation) (i) how do these intuitions formally manifest in biological networks? (ii) what are the possible signature structures and very importantly (iii) whether a generalised result can be obtained for networks of any size? These are somewhat formidable questions, especially given the non-linear nature of biological processes. However, it turns out that linear systems theory can still provide concrete answers. Essentially, the linearized structure of the system provides the answer to a binary question of whether the network is able to provide adaptation or not. If yes, further conditions on the linearized system are obtained and the problem of determining suitable network structure is resolved. The proposed framework is systematic and generic as against computationally demanding search methods and finding specific control strategies for a particular network to achieve adaptation.

Deriving the necessary conditions for adaptation, we show that a minimum of N edges are required for an N-node network to produce adaptation. We use this result to deduce further, that there exist only two ways, namely (1) feedback loop, and (2) multiple forward paths in an N−node network, to provide adaptation.

Finally capturing the above results in Theorem 6, we show that existence of either a negative feedback loop or incoherent feed forward node acts as a necessary condition for adaptation. This result agrees with the observations in the seminal work of Tang and coworkers [10], but without the need for elaborate simulations and parameter samplings. We believe that the conditions obtained for a general N-node network assumes most prominence for two reasons: (i) no prior theoretical results exist and (ii) it provides a deeper understanding of how a general protein network is configured to provide adaptation. Lemma 7 establishes that adaptation is retained in presence of a canonical downstream connection. This non-retroactive nature of these networks implies that they are highly likely to preserve their function in synthetic circuits designed with various modules.

It should also be noted that the topologies obtained from the linearized hyperbolic system provide perfect adaptation in the practical (nonlinear) scenario. The more generic case comprising of the possibility of a non-hyperbolic system providing adaptation can be an interesting future study. Also, the controllability condition used in this paper works as a sufficient condition for the controllability of the actual nonlinear system. The area of nonlinear controllability can be explored in this context to avoid missing out on false negatives.

In sum, we see four definitive contributions of this study. We first proved via Theorem 1 that the network structures for adaptation ipso facto reduce peak time because of the infinite precision (zero-gain) requirement. Second, the question of adaptation for staircase-type disturbances had been addressed and concise conditions inspired from systems theory were proposed regarding this for the first time. Third, we argue that the structural conditions obtained as the necessary conditions for adaptation herein, are most stringent among the ones in the existing literature (Refer to Table S1 in the *SI*). Araujo *et al.* (2018) emphasised the need of either a loop or multiple opposing forward paths whereas this paper extends this result further arguing that the sign of at least one feed back loop has to be negative for ensuring adaptation in absence of opposing forward paths [30]. Fourth and most notably, the entire algorithm remains agnostic to the particularities of the reaction kinetics. Our approach lays the foundation for the application of LTI systems theory to predict topologies and fine-grained constraints, for networks capable of achieving other functionalities.

## Acknowledgement

The authors thank K R Ghusinga for valuable comments on the manuscript. PB acknowledges funding from the Ministry of Education, Government of India.

## Supporting Information

This section presents the necessary calculations, proofs and the rate laws used for simulation studies.

### 1 Two-node networks

Considering the inability of a single protein network to provide adaptation, we now turn to a two-protein network. The network comprises two proteins 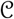 and 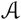, which are connected; 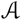 is further connected to the external source of disturbance (input 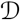), and the concentration of 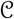 is considered as the “output species”. Let us denote the concentration of 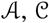 and the disturbance species 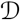 by x_1_(t), x_2_(t),and d(t) respectively. The resultant linearized state space representation is:

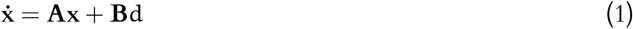

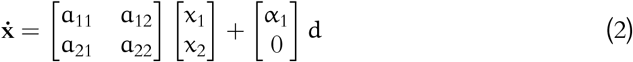

According to the previously derived conditions for adaptation (Eqs. (9) and (5)), the output state x_2_ has to be controllable by the applied input. This demands a non-zero value for a_21_, *i. e.* there should exist an edge from 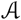 to 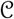. As per the second condition for adaptation, the final value of the linearized output state x_2_ should be zero, and the system Matrix **A** should be Hurwitz.

Denote the steady-state value as 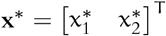. Then, at steady state,

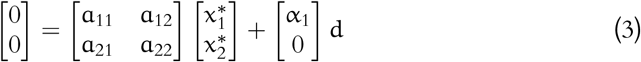

For any vector of the form 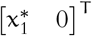 to be a solution to the above system of equations requires a_21_ to be zero. This is a violation of the controllability condition. Therefore, it can be concluded that a two-node network with different input–output nodes *cannot provide adaptation*.

To examine an alternate possibility, let us now consider the input node 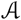 itself as the output node as well. Note that the state x_1_ is always controllable by the disturbance ∀α_1_ ≠ 1 Also, if a_22_ is made zero possibly with a positive self loop on 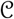, then, the final steady-state value of x_1_ can be zero, irrespective of x_2_. In this case, for **A** to be a stable, a_21_a_12_ has to be negative. This condition maps to a negative feedback between 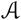 and 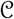 (Figure S1). Taken together, the admissible topology must have

1. a_22_ = 0, ⟹ possible positive self loop on 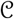
2. a_21_a_12_ < 0 ⟹ negative feedback between 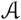 and 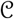.

#### 1.1 Toilet Flush Phenomenon

To demonstrate further, let us consider a network of three proteins, 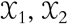, and 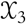, where 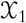 is connected with 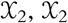 is connected with 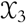, and 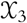 is connected with 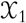. Let the output node, 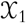, be perturbed with an input, u. If we adopt mass-action kinetics and assume the total mass to be conserved, *i. e.* 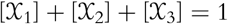, thereby leaving two independent states, the state equation can be written as

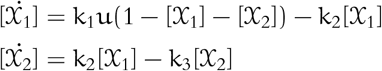

For the case of zero input, the steady-state values are 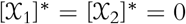. It can be shown by our method that, for adaptation, k_3_ has to be zero, but after a single step, the steady-state values of the states become [0, 1], thereby rendering the system linearized around the new steady-state uncontrollable rendering the system responsive only for the first step type jerk.

**Fig S1.**
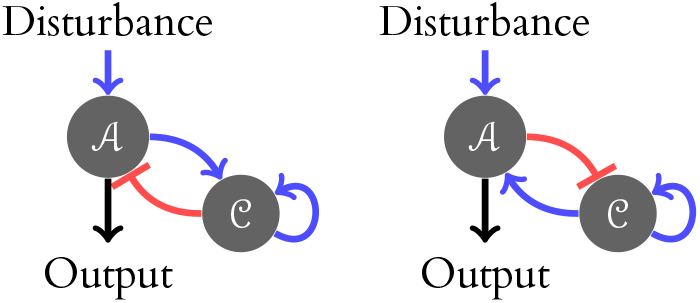
Admissible two-node topologies. The normal (blue) arrowheads signify activation, while the bar-headed (red) arrows signify repression.

## 2 Equivalence between conditions between adaptation

It was shown in the previous literature that the condition for adaptation is 1) one of the zeros in the transfer function to be placed in the origin. 2) In this work, we have shown for a system (**A**, **B**, **C**, **D**) to provide adaptation the necessary condition is **CA**^−1^**B** = 0. We argue that these two claims are equivalent. To prove this claim, we first establish 1 → 2.

*Proof* A proper and stable transfer function H(s) which provides adaptation can be expressed as

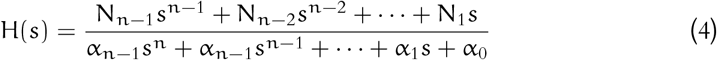

The corresponding state space representation (**A**, **B**, **C**, **D**) can be written assuming zero pole zero cancellation (full controllbility) can be obtained as

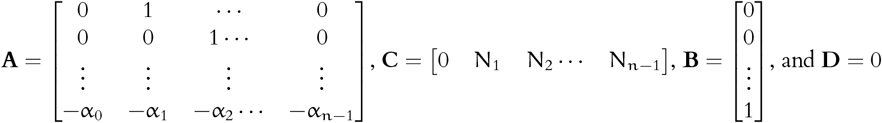

With the structure of (**A**, **B**, **C**, **D**) it can be seen that **CA**^−1^**B** = 0 which proves the forward assertion.

Subsequently, it is to be proved that the zero at origo condition amounts to the condition derived in the main script.

*Proof* : For a given state space structure (**A**, **B**, **C**, **D**) the transfer function can be written as

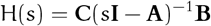

. The zero at the origo means zero final value of the step response (Y(s)) of the system.

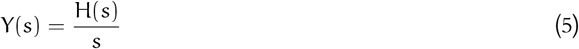

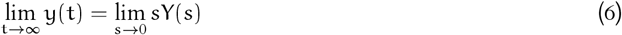

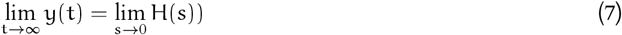

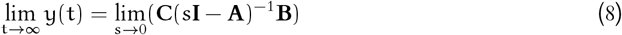

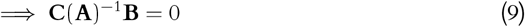

So, it can be seen that both the assertion and its converse are true so the condition for adaptation derived in this work is equivalent to the standard condition of zero at the origo.

### 2.1 Derivation of (23) from (5)

: In this subsection, we argue that the infinite precision condition derived in equation (5) is a more general than the one derived in (23). The infinite precision condition is obtained as

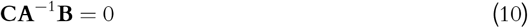

For the specific case of (23), the input disturbance is applied on the first node and the output is considered as the concentration of the k^th^ node. Therefore the **B** ∈ ℝ^N^ and **C** ∈ ℝ^1×N^ matrix are of the form β**e**_1_ and 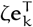 respectively where **e** ∈ ℝ^N^ are unit vectors across the j^th^ axis and β, ζ are nonzero scalars. It is to be noted that since **A** is Hurwitz as per the stability condition the determinant is non-zero and will be the denominator in the expression of **CA**^−1^**B**. According to the very definition of matrix inverse we know that the (i, j)^th^ element of det(**A**)**A**^−1^ refers to the minor of the (j, i)^th^ component of **A**. Due to the specific structure of **B**, **A**^−1^**B** will be a scaled version of the first column of **A**^−1^. Similarly with the given structure of **C** the expression **CA**^−1^**B** returns a scaled version of the (k, 1)^th^ element of det(**A**)**A**^−1^ which in turn is the minor of the (1, k)^th^ component of the **A** matrix.

## 3 Generalization

This section deals with the necessary results and demonstrations that act as the stepping stones for the results shown in the main text.

### 3.1 Two principal means of achieving infinite precision

: The infinite precision equation represented in (23) involves computation of the minor of the term that maps back to an edge from the output to the input node. In an N−node network (x_1_, X − 2, … , x_N_ as the concentration of the 1^st^, 2^nd^, … , N^th^ node respectively), if the concentration of the input node is considered as the first node (concentration x_1_) and the k^th^ node as output with the respective concentration expressed as x_k_, then according to (23), the

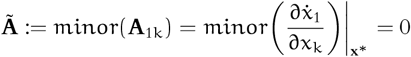

, For the system matrix **A** ∈ ℝ^N×N^ there are N! number of terms present in the determinant expression in which 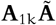 involves (N − 1)! number of terms. From combinatorial matrix theory, it is well known that ([35]) each term in the determinant expression of any diagraph matrix can be expressed as the product of the diagonal entries and loops with no mutual nodes. Following this, it can be said that each term of 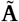 contains exactly one forward path from the input to the output node. It is to be noted that each of the (N − 1)! terms in the expression of 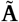 contains N − 1 elements. In terms which refer to the forward paths with less than N − 1 number of edges, the remaining entries are composed of the diagonal and the loop elements. It is obvious that there are two ways in which all the terms of 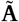 sum up to zero

1. All the terms are zero individually.
2. There exist terms with equal and opposing actions.

As discussed earlier 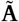 contains (N − 1)! terms. Each term contains exactly one forward path from the input to the output node. One option can be to have a network without any forward path but this leads to uncontrollability of the output node. So the only other option is to make all the forward paths with N − 1 edges absent along with at least one of of the diagonal elements to be zero such that all the terms are individually zero. This is exactly what is referred as the opposer module in [30] In the second case, the non-zero terms can be grouped in to three classes. In this context, let us define certain notations and functions that shall be helpful in putting things in perspectives. Suppose set ℕ_PL_ contains all the forward paths and loops of the network, set 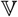 contains all the nodes. Also, 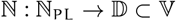 returns all the nodes involved in a given forward path 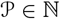. Then, cardinality of the set 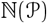 provides the number of nodes involved in the forward path 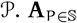 refers to the component of the **A** matrix that represents P. Since, **A** acts as the diagraph matrix this is a ono-to-one mapping. Therefore, P and **A**_P_ shall be used interchangeably to reduce the abundant notations. i) Let us consider two forward paths F_1_ ∈ ℕ_PL_ with f_1_ nodes, F_2_ ∈ ℕ_PL_ with f_2_ nodes and a loop L involving p nodes such that 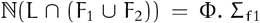 is the permutation set of all diagonals (N − f_1_ − p − 1) except the ones situated in 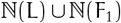 similar notations are also invoked for F_2_. For this case, the expression of 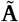 concerning the aforementioned loops. forward paths can be written as

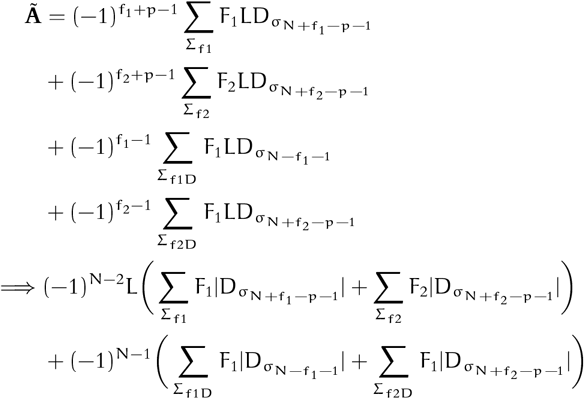

Now, the only way to achieve 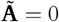 while ensuring stability ((35)) is to have sign(F_1_) = (−1)sign(F_2_) ii) Let us consider two forward paths F_1_ ∈ ℕ_PL_ with f_1_ nodes, F_2_ ∈ ℕ_PL_ with f_2_ nodes and two loops L_1_, L_2_ involving p_1_ and p_2_ nodes such that ℕ(L_1_) ∩ ℕ(F_2_) = N_j_, ℕ(L_2_) ∩ ℕ(F_1_) = N_k_ and ℕ(L_1_) ∩ ℕ(L_2_) = N_l_. It is to be noted that in this case, apart from F_1_ and F_2_ there exist two other forward paths 1) From the input node (denote as node 1) to the 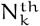 node via F_1_, then from 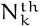 to the 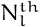 node via L_2_ and lastly from 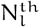 to the 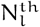 via L_1_ and from 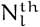 to output node (denote as k^th^ node) via F_2_. Let us call this as F_12_ 2) From the input node (denote as node 1) to the 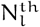 node via F_2_, then from 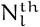 to the 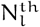 node via L_1_ and lastly from 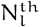 to the 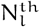 via L_2_ and from 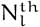 to output node via F_1_. Let us denote this as F_21_ In this case as well the terms in the expression of 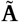 shall be similar to the previous case except an addition of two forward paths F_1_ and F_2_. Now, the only way to mutually cancel the terms in 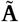 concerning the forward path F_1_ and F_2_, assuming F_1_ and F_2_ are of the same sign is to have sgn(L_1_) = (−1)sgn(L_2_) in that case it can be seen that sgn(F_12_F_21_) = sgn(L_1_L_2_) = −1. This means the forward paths F_12_ and F_21_ are of the opposite sign. iii) Let us consider two forward paths F_1_ ∈ ℕ_PL_ with f_1_ nodes, F_2_ ∈ ℕ_PL_ with f_2_ nodes and two loops L_1_, L_2_ involving p_1_ and p_2_ nodes such that ℕ(L_1_) ∩ ℕ(F_2_) = N_j_ , ℕ(L_2_) ∩ ℕ(F_1_) = ℕ_k_ and ℕ(L_1_) ∩ ℕ(L_2_) = Φ. The corresponding expression for 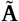 can be written as

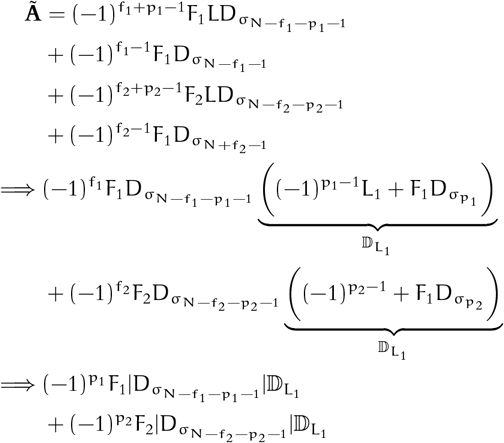

Assume F_1_ and F_2_ are of the same sign then

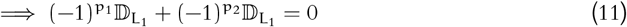

Again, for stability, we know

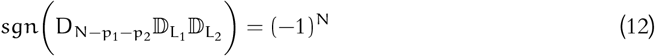

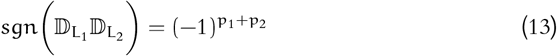

The only way to satisfy (11) 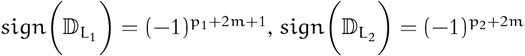 or 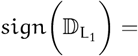 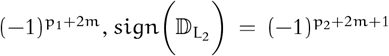 where, 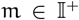. In both the cases 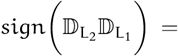 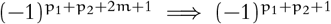

This again is the violation of the stability condition depicted in (13). Therefore the only way to drive 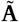 to zero is to have incoherrent feedforward paths considering all the diagonal elements are non-zero and negative.

It has already been established in the main text that in order for the network to be able to provide adaptation, it has to be controllable with respect to the external disturbance. In the following theorem, we argue that there exists at least one forward path from the input to the output node for the system to be controllable.

#### 1. Theorem

*For an* N−*node network with different input and output nodes, considering the states as the concentration of the proteins the resultant state space system is output controllable if there exists at least one forward path from the input to the output node.*

*Proof.* In order to prove the above theorem, we have to show that the system is not output controllable if there exists no forward path from the input node to the output node.

Without any loss of generality, let us denote the input node as the first node with concentration x_1_ and the same for the k^th^ node (x_k_) is considered as the output. Assume, there are p nodes which are connected with the input node in such a way that there exists at least one forward path from the input node to all of the P nodes. None of the remaining N − P nodes can be reached from the input node. Using the property that the system matrix **A** for the linearised state space system acts as a digraph matrix for the network,

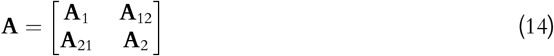

where, **A**_1_ ∈ ℝ^P×P^ captures the inter connections among the P nodes reachable from the input node, **A**_12_ ∈ ℝ^P×N−P^ contains the connections from the N − P nodes to the first P nodes, **A**_12_ ∈ ℝ^N−p×P^ contains the connections from the first P nodes to the remaining N − P nodes, and **A**_2_ ∈ ℝ^N−P×N−P^ reflects the interconnections among the last N − P nodes. Since there exists no froward path from the input node to any of the N − P nodes **A**_21_ is a zero matrix. The actuator matrix **B** can be written as

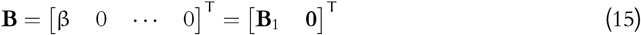

where, **B**_1_ ∈ ℝ^1×P^ is an elementary vector with the first element being non-zero (β) as the input node is considered as the first node. Given the pair (**A**, **B**) the controllability matrix (Γ_c_) can be written as

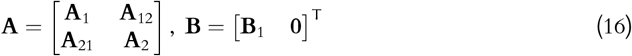

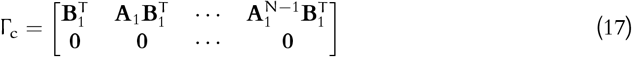

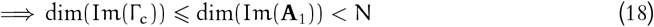

where Im(.) denotes the column space of a matrix and dim(.) calculates the dimension of a given vector space. From (18) it is clear that the Kalman rank condition can not be achieved in this case.

### 3.2 Hurwitz stability of Negative feedback loops and IFFLP

The Hurwitz condition in systems theory guarantees the exponential asymptotic stability of the linearised system. Further, the Hartman-Grobman theorem ensures the stability of the corresponding non-linear system if its linearised counter part is exponentially stable. Therefore, to comment on the stability of the actual non linear system, we first investigate whether the system matrix **A** of the linearised system is Hurwitz. For any matrix **A** ∈ ℝ^N×N^ to be Hurwitz, one of the necessary conditions is the following

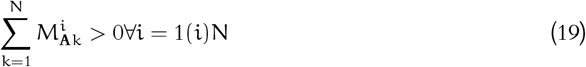

where, 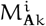 is the all possible i^th^ principal minors of **A**. It is evident from (19), there are N conditions that need to be satisfied for any N×N matrix to be Hurwitz. As established before, the linearised system matrix **A** can be considered as the diagraph matrix of the associated network structure. In this scenario, the sum of all possible i^th^ principal minors can be expressed as all possible i−node loops present in the network structure, loops with less than i nodes and diagonals. To illustrate further, assume the network has two loops L_1_ and L_2_ containing N_1_ and N_2_ number of nodes. Further, assume there exists no common nodes in L_1_ and L_2_. In that case, the expression for the sum of all i^th^ (i > N_1_ + N_2_) principal minor can be written as

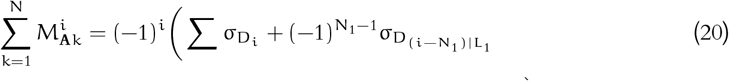

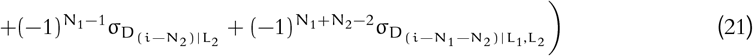

where, σ_i|t_ is the permutation operator that chooses k diagonals from the set of N (**A** is N × N) diagonal elements, the subscript t means the choice of i diagonal elements should be such that it does not have any common co-ordinate with the elements in t. For a network with a single loop (L_p_) of p_1_ nodes and cumulative sign being negative the sum of all the principal minors of order i can be written as

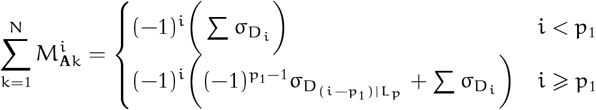

It can be seen in both the scenarios (i < p_1_, i ⩾ p_1_) the sign of the sum of i^th^ order minor is always positive given the diagonals and the L_1_ is of negative sign. Hence presence of negative feedback loop satisfies the Hurwitz condition for exponential stability.

In the case of feedforward networks without any loop the sum of the i^th^ principal minors shall always be sum of the combination of ∀i diagonal elements chosen from N diagonals in which case, the sum of the principal minors shall always be positive i given the diagonal elements are negative. This also guarantees the Hurwitz property of the networks with only feedforward paths.

For an N × N matrix there are N! number of terms present in the determinant expression. It can be proved that every term in the expression contains at least one loop except the product term of the diagonals. The elements which carried a single loop were discussed in the main text and it was shown that the elementary motif associated with one of these terms need to be of negative feedback type i,e the loop sign should be negative. The elements containing multiple non-overlapping loops can not provide adaptation for the associated network becomes uncontrollable. For these networks it can be shown that if the cumulative sign of all the loops are positive then also it can satisfy the determinant condition i,e, the sign of the determinant becomes (−1)^N^. These networks along with another link/loop (to make the network controllable) leads to Hurwitz instability by making at least one of eigenvalues positive. Following is an illustration of a four node network. Assume a five node network which has two loops one involving 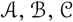 and the other with 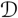 and 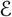. The concentration states of 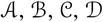 are represented as x_1_, x_2_, x_3_, x_4_, x_5_ respectively. Input (I) is applied on 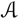 and the concentration of 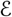 is considered as output.

From the network structure in S2, it can be seen that there are two loops involved in the network. One is engaging 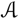 and 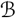, another with 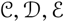 nodes. Both the feedback schemes are positive in nature. From the structure it can be intuitively seen that the network is controllable for any non-zero strength of the edge from 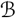 to 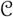. This can also be proved mathematically by evaluating the rank of the associated controllability matrix.

**Fig S2.**
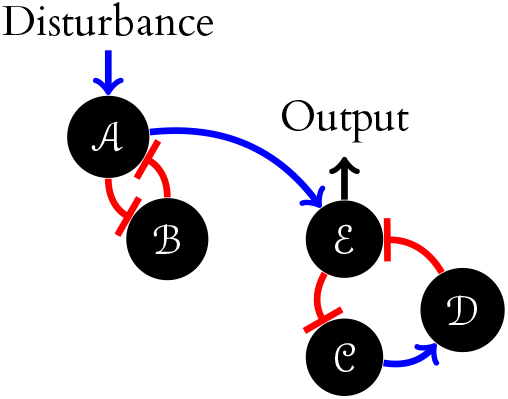
Proposed five node network which can not provide adaptation albeit satisfying the weaker condition for stability

For this network to provide adaptation, the corresponding system matrix **A** after linearisation can be of the structure

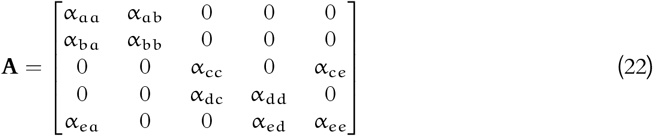

Note, if there is no edge from 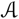 to 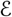, the network would be uncontrollable. The condition to be met for this five node network to provide adaptation is the following 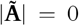, where 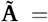 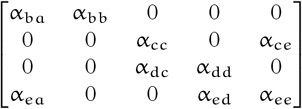 So, for 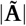 to be zero α_cc_ has to be zero. The next condition is concerning the stability of **A**. With α_bb_ = 0 the determinant of **A** can be written as

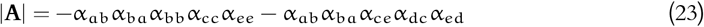

Now, for the system to be Hurwitz stable, the determinant of **A** is necessarily of the sign (−1)^5^=−1. This can be achieved in two ways 1) both the terms are negative or 2) Either one of them is negative with magnitude greater than that of the positive term. The first case leads to at least one negative feedback, preferably between 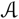 and 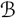. In the second case, if both the loops are of positive feedback and if

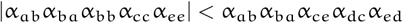

then the necessary condition for the Hurwitz stability of **A** is satisfied. But on a careful introspection, it can be seen that at least one of the eigenvalues of **A** is positive which goes to violate the Hurwitz stability condition for **A** thereby leading to instability. So, the above network structure can be ruled out.

This can be understood from the **A** matrix for these cases. To make the network controllable and able to provide adaptation, it is necessary to add an edge from the input to the output node. Although the addition of an element changes the spectrum of the overall matrix, the spectrum of the block matrices containing the loops other except one will not be changed. If all the loops are positive at least one of the eigenvalues of the block matrices will be positive leading to instability for the overall matrix. In the example of S2 the addition of an edge from 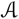 to 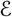 has changed the spectrum of **A** without changing the spectra of the block matrix 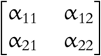. With 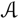 and 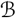 in positive feedback, one of the eigenvalues can be verified as positive, which leads to the violation of Hurwitz property of **A**.

**Table S1.**
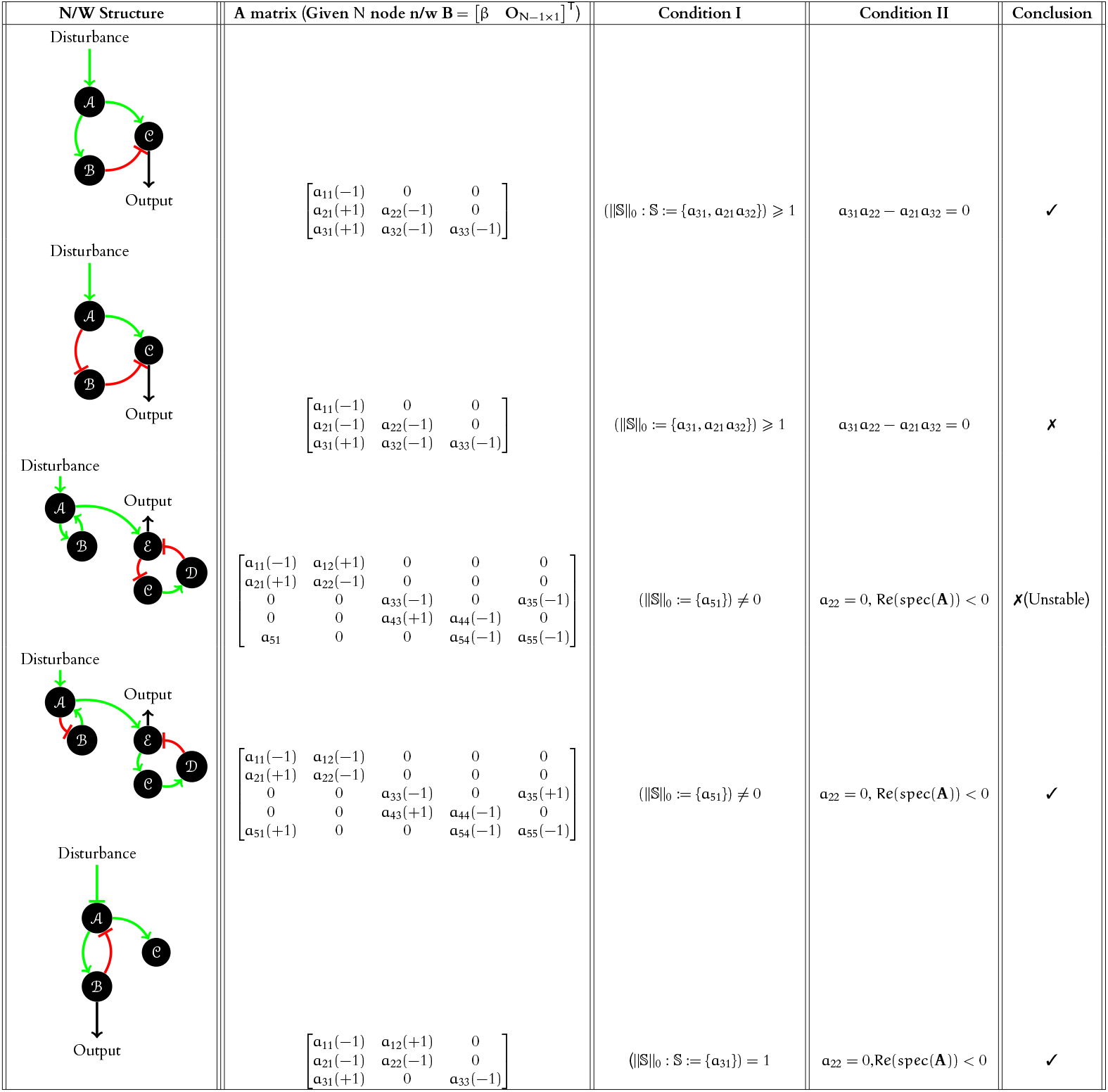
Demonstration of the algorithm. 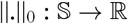 refers to the number of non-zero elements in the set 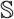.

## 4 Equations for simulation

### Two node network

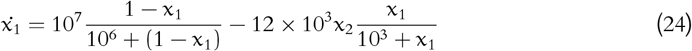

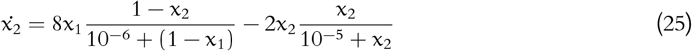

### Voltage gated Na ion channel

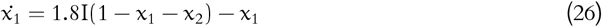

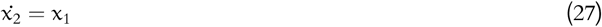

### IFFLP

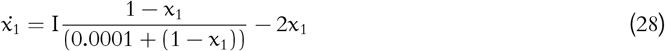

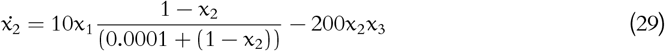

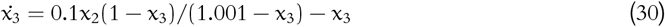

### NFBLB

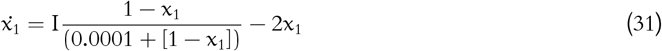

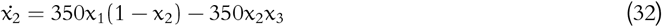

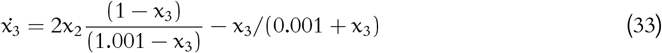

### IFFLP+NF

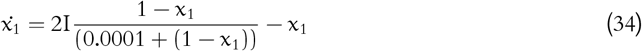

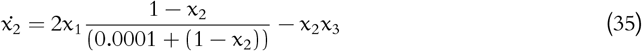

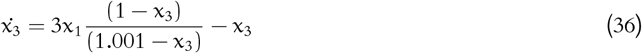

### IFFLP+NF

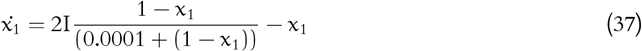

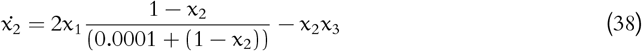

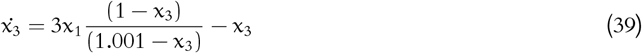

## References

1. Voit E. A First Course in Systems Biology. 1st ed. Garland Science; 2012.

2. Raman K. An Introduction to Computational Systems Biology: Systems-Level Modelling of Cellular Networks. 1st ed. Boca Raton, FL: Chapman and Hall/CRC; 2021.

3. Hans D, Arunn V, Holden L, Folke O. Chaos in biological systems. Springer US; 1987.

4. Kulkarni V, Stan G, Raman K. A Systems Theoretic Approach to Systems and Synthetic Biology I: Models and System Characterizations. Springer Netherlands; 2014.

5. Milo R, Shen-Orr S, Itzkovitz S, Kashtan N, Chklovskii D, Alon U. Network motifs: simple building blocks of complex networks. Science. 2002;298(5):824–827.

6. Constantino P, Tang W, Daoutidis P. Topology Effects on Sparse Control of Complex Networks with Laplacian Dynamics. Scientific Reports. 2019;9(9034):1–9. doi:10.1038/s41598-019-45476-6.

7. Wang Z, Su Q, Huang G, Wang X, Wang W, Grebogi C, et al. A geometrical approach to control and controllability of nonlinear dynamical networks. Nature Communications. 2016;7(11323). doi:10.1038/ncomms11323.

8. Liu Yang Y, Slotine J, Barabási A. Controllability of complex networks. Nature. 2011;473(167):167–173. doi:https://doi.org/10.1038/nature10011.

9. Golubitsky M, Wang Y. Infinitesimal homeostasis in three-node input-output networks. Journal of Mathematical Biology. 2020;80(4):1163–1185.

10. Ma W, Trusina A, El-Samad H, Lim A, Tang C. Defining Network Topologies that Can Achieve Biochemical Adaptation. Cell. 2009;138(4):760–773.

11. Königs V, de Oliveira Freitas Machado C, Arnold B, Blümel N, Solovyeva A, Löbbert S, et al. SRSF7 maintains its homeostasis through the expression of Split-ORFs and nuclear body assembly. Nature Structural and Molecular Biology. 2020;27(3):260–273.

12. Bernardo M, Yuhai T. Perfect and near-perfect adaptation in a model of bacterial chemotaxis. Biophysical Journal. 2003;84(8):2943–2956.

13. Tyson J. On the Existence of Oscillatory Solutions in Negative Feedback Cellular Control Processes. Journal of Mathematical Biology. 1975;1:311–315. doi:https://link.springer.com/content/pdf/10.1007%2FBF00279849.pdf.

14. Li Z, Liu S, Yang Q. Incoherent Inputs Enhance the Robustness of Biological Oscillators. Cell. 2017;5(12):72–81.

15. Ananthasubramaniam B, Herzel H. Positive Feedback Promotes Oscillations in Negative Feedback Loops. PLoS One. 2014;9(8):1–11. doi:pone.01047611..11.

16. Novak B, Tyson J. Design principles of biochemical oscillators. Nature Reviews Molecular Cell Biology. 2008;9(12):981–991.

17. Angeli D, Ferrell J, Sontag E. Detection of multistability, bifurcations, and hysteresis in a large class of biological positive-feedback systems. Proceedings of the National Academy of Sciences USA. 2004;101(7):1822–1827. doi:10.1073/pnas.0308265100.

18. Torday J. Homeostasis as the Mechanism of Evolution. Biology (Basel). 2015;3(3):573–590.

19. Friedlander T, Brenner N. Adaptive response by state-dependent inactivation. Proceedings of the National Academy of Sciences USA. 2009;106(11):22558–22563.

20. Briat C, Gupta A, Khammash M. Antithetic Integral Feedback Ensures Robust Perfect Adaptation in Noisy Biomolecular Networks. Cell Systems. 2016;2(10):15–26.

21. Ferell J. Perfect and Near-Perfect Adaptation in Cell Signaling. Cell Systems. 2016;2(7):62–67.

22. Sontag E. Adaptation and regulation with signal detection implies internal model. Syst and Cont letters. 2003;50(16):119–126. doi:10.1016/S0167-6911(03)00136-1.

23. Waldherr S, Streif S, Allgöwer F. Design of biomolecular network modifications to achieve adaptation. IET Syst Biol. 2012;6(14):223–31. doi:10.1049/iet-syb.2011.0058.

24. Drengstig T, Ueda H, Ruoff P. Predicting perfect adaptation motifs in reaction kinetic networks. J Phys Chem B. 2008;112(15):16752–16758. doi:10.1021/jp806818c.

25. Drengstig T, Kjosmoen T, Ruoff P. On the Relationship between Sensitivity Coefficients and Transfer Functions of Reaction. J Phys Chem B. 2011;115(16):6272–6278. doi:10.1021/jp200578e.

26. Bhattacharya P, Raman K, Tangirala A. A systems-theoretic approach towards designing biological networks for perfect adaptation. IFACPapersOnline. 2018;51(5):307–312. doi:10.1016/j.ifacol.2018.05.033.

27. Yi T, Huang Y, Simon M, Doyle J. Robust perfect adaptation in bacterial chemotaxis through integral feedback control. Proceedings of the National Academy of Sciences USA. 2000;97(9):4649–4653. doi:10.1073/pnas.97.9.4649.

28. Marcelo B, Nan H, Dohlman G, Timothy C. Mathematical and Computational Analysis of Adaptation via Feedback Inhibition in Signal Transduction Pathways. Biophysical Journal. 2007;93(3):806–821.

29. Jamal S, Rahi, Johannes L, Kresti P, Alexander K, N M, et al. Oscillatory stimuli differentiate adapting circuit topologies. Nature Methods. 2017;14(10):1010–1016.

30. Robyn A, Lance L. The topological requirements for robust perfect adaptation in networks of any size. Nature Communications. 2018;9(13):1757–1769.

31. Del Vecchio D. A control theoretic framework for modular analysis and design of bio-molecular networks. Annual Reviews in Control. 2013;7(6):333–345.

32. Hespanha Joao P. Linear Systems Theory: Second Edition. Princeton University Press; 2018.

33. Bhattacharya P, Raman K, Tangirala A. Systems-Theoretic Approaches to Design Biological Networks with Desired Functionalities. Methods in Molecular Biology. 2021;2189:133–155.

34. Goh LK, Sorkin A. Endocytosis of Receptor Tyrosine Kinases. Cold Spring Harb, Perspective Biology. 2013;5(14):833–849. doi:10.1101/cshperspect.a017459.

35. Maybee J, Driessche P, Olesky D, Wiener G. Matrices, Digraphs, and Determinants. Society of Industrial and Applied Mathematics. 1989;10(4):500–519.

